# The membrane insertion of the pro-apoptotic protein Bax is a Tom22-dependent multi-step process: a study in nanodiscs

**DOI:** 10.1101/2024.02.16.580668

**Authors:** Akandé Rouchidane Eyitayo, Laetitia Daury, Muriel Priault, Stéphen Manon

## Abstract

Membrane insertion of the pro-apoptotic protein Bax was investigated by setting up cell-free synthesis of full-length Bax in the presence of pre-formed nanodiscs. While Bax was not spontaneously inserted in nanodiscs, co-synthesis with the mitochondrial receptor Tom22 promoted Bax membrane insertion. The initial interaction of Bax with the lipid bilayer exposed the hydrophobic GALLL motif in Hα1 leading to Bax precipitation through hydrophobic interactions. The same motif was recognized by Tom22, triggering conformational changes leading to the extrusion and the ensuing membrane insertion of the C-terminal hydrophobic Hα9. Tom22 was also required for Bax-membrane insertion after Bax was activated either by BH3-activators or by its release from Bcl-xL by WEHI-539. The effect of Tom22 was impaired by D^154^Y substitution in Bax-Hα7 and T^174^P substitution in Bax-Hα9, that are found in several tumors. Conversely, a R^9^E substitution promoted the spontaneous insertion of Bax in nanodiscs, in the absence of Tom22. Both Tom22-activated Bax and BaxR ^9^E alone permeabilized liposomes to dextran-10kDa and formed ∼5nm-diameter pores in nanodiscs. The concerted regulation of Bax membrane insertion by Tom22 and BH3-activators is discussed.

## Introduction

The mitochondrial apoptotic pathway involves proteins of the Bcl-2 family, including anti-apoptotic proteins, such as Bcl-2 and Bcl-xL, multi-domain pro-apoptotic proteins Bax, Bak and Bok, and pro-apoptotic BH3-only proteins, such as Bid and Bim ([1,2,] for recent reviews). In healthy cells, Bax is inactive in the cytosol or loosely bound to intracellular membranes. It becomes active following apoptotic signals that induce a series of events culminating in the formation of a pore that permeabilizes the mitochondrial outer membrane (MOM) to apoptogenic factors.

Soluble, inactive Bax is structured in 9 α-helices, organized around a hairpin formed by Hα5 and Hα6 [3]. The C-terminal Hα9 is hydrophobic, like the C-terminal membrane anchors of anti-apoptotic proteins Bcl-2 and Bcl-xL [4,5]. However, Bax-Hα9 does not drive the MOM-insertion of Bcl-xL while the C-terminal α-helix of Bcl-xL drives the MOM-insertion of Bax [3,6]. Indeed, Bax Hα9 is stabilized in a hydrophobic groove formed by the BH domains, owing to the presence of hydrophobic residues on its core facing side and hydroxyl residues on its outside facing side [3]. The only hydroxyl residue on the core facing side, S^184^, is stabilized by a hydrogen bond with D^98^ in Hα4. The deletion of S^184^, or its substitution by a non-hydroxylated residue, promotes the spontaneous insertion of Bax in both reconstituted and cellular models [3,7]. Hα9 extrusion from the hydrophobic groove thus appears as a crucial step towards Bax membrane-insertion. The BH3-only protein Bim binds at a site close to Hα1, α1-α2 loop and Hα6 (the so-called “non-canonical” binding site) [8], resulting in long-range conformational changes that may ultimately lead to the extrusion and the insertion of Hα9 [9]. Accordingly, mutagenesis-induced conformational changes in the α1-α2 loop prevented Hα9 extrusion [10]. Anti-apoptotic proteins competing with Bim-binding might also block these long-range conformational changes [11].

The N-terminal extremity of Bax is also involved in the regulation of its mitochondrial localization. Indeed, the absence of the 19 N-terminal residues generates a variant, called BaxΨ, having a constitutive mitochondrial localization [12], which was found in low grade glioblastomas displaying high level of apoptosis [13]. Bax-Hα1 (residues 16-35) fused to RFP competed with the mitochondrial addressing of BaxΨ suggesting that a receptor for Hα1 was involved in Bax mitochondrial localization [14]. Hα1 was further shown to interact with the mitochondrial receptor Tom22, a component of the TOM complex (Translocase of Outer Membrane) [15]. Tom22 down-regulation decreased Bax mitochondrial localization in apoptotic mammalian cells [15,16] and following heterologous expression in yeast [15,17] and in *Drosophila* [18]. However, because other studies did not report any significant role of TOM components on Bax localization in mammalian cells [19] nor in yeast [20], the potential role of Tom22 in Bax mitochondrial localization remains debated. Another TOM subunit, namely Tom20, is involved in the mitochondrial localization of Bcl-2 [21,22] and Metaxins 1 and 2, components of the SAM complex (Sorting and Assembly Machinery), are involved in the mitochondrial localization and activation of Bak [23,24]. It therefore appears that the role of MOM receptors in the targeting, activation and regulation of Bcl-2 family members deserves further investigations ([25,26] for reviews).

The reconstitution of membrane proteins in nanodiscs is a successful approach not only to resolve the structure of membrane proteins but also to gain insight into mechanistic aspects ([27–29] for reviews). A cryoEM study of Bax activated by Bid-BH3 at low pH, and inserted into nanodiscs, showed the occurrence of a small pore (∼3.5nm in diameter) [30]. On this basis, molecular dynamics simulations supported that the reorganization of a limited number of lipid molecules around Bax could generate a toroidal pore also involving Hα5 and Hα6 [31]. We have set-up a more direct approach by running the cell-free synthesis of Bax in the presence of pre-formed nanodiscs [32]. We used full-length BaxWT and a mutant BaxP^168^A, that was previously shown to be spontaneously membrane-inserted in glioblastoma cells [33], yeast cells [34] and liposomes [35]. However, both variants largely precipitated in the presence of nanodiscs and therefore failed to be inserted [32]. The precipitated fraction, containing pure Bax, was used to co-form nanodiscs: interestingly, TEM studies showed that nanodiscs containing BaxP^168^A had a larger size than those containing BaxWT, and that some of them displayed a pore having a diameter close to 5nm [32]. However, the objects were heterogeneous, precluding structural investigations. Subsequently, we observed that the cell-free co-synthesis of Bax with its anti-apoptotic partner Bcl-xL induced the spontaneous co-insertion of both proteins into nanodiscs [36].

These data urged us to further investigate the mechanisms underlying Bax interaction with nanodiscs and the possible role of Tom22, by setting-up a minimal system involving the co-synthesis of Bax and Tom22 in the presence of nanodiscs. By using several Bax mutants, we showed that Tom22 interacted with Bax Hα1 to promote a conformational change resulting in the extrusion and the membrane insertion of Bax Hα9. Tom22 was also required for Bax-membrane insertion following Bax activation by BH3-activators or by its interaction with Bcl-xL then release by WEHI-539. Furthermore, Tom22 promoted the ability of Bax to form a pore. We propose a model that emphasizes the concerted action of Tom22 and BH3-activators in Bax targeting, insertion and pore formation.

## Results

### The co-synthesis of Tom22 prevents Bax precipitation and stimulates Bax insertion into nanodiscs

As reported previously [32], the presence of nanodiscs induced the precipitation of a large fraction of Bax (Fig.S1), while nanodiscs remained in the supernatant (Fig.1A). The fraction of Bax that was not precipitated still failed to interact with nanodiscs, as shown by its detection in the unbound fraction of Ni-NTA-sepharose affinity chromatography to immobilize the His7-tagged nanodisc scaffold protein MSP1E3D1 (Fig.1C).

**Figure 1:**
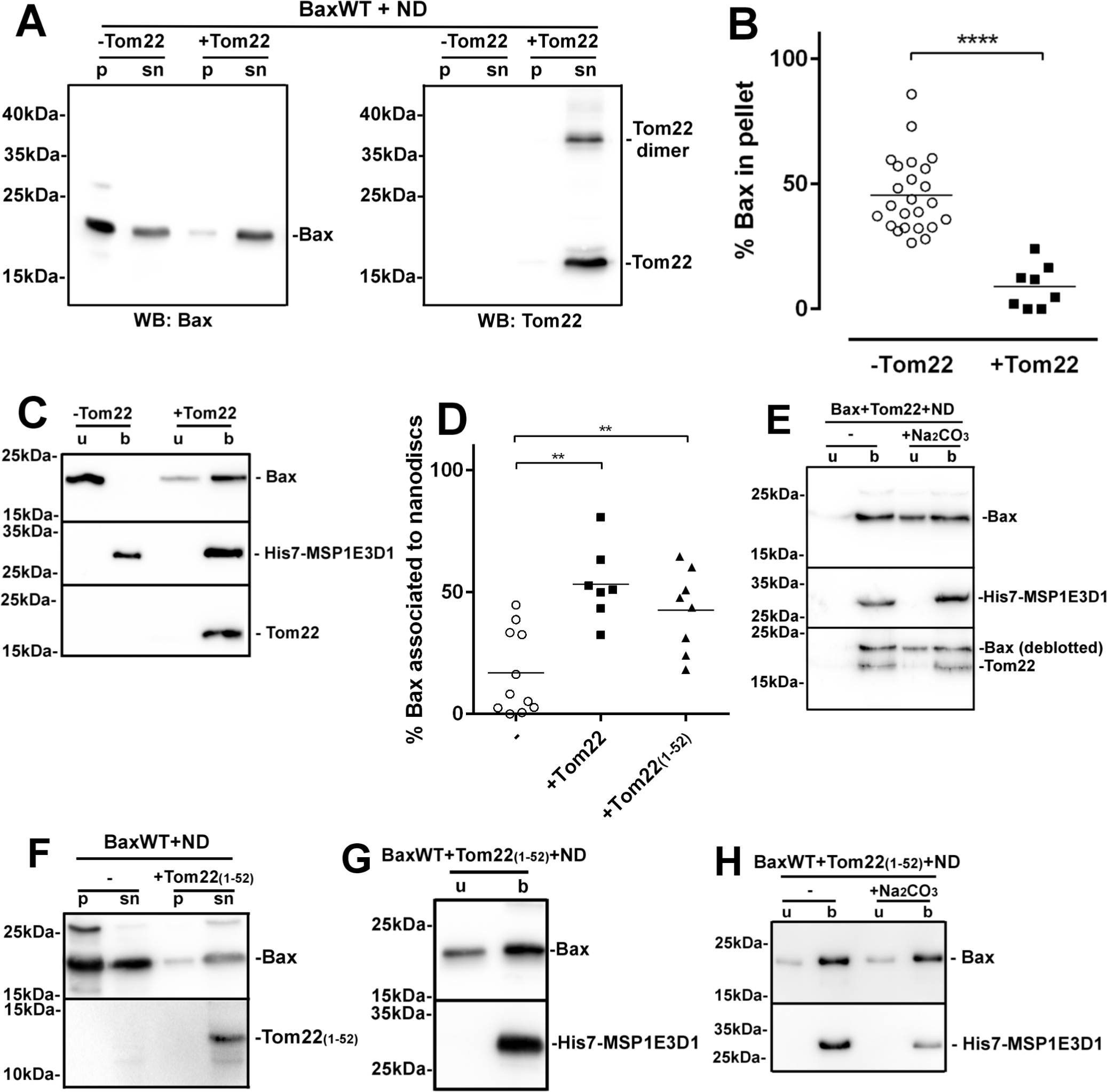
Tom22 co-synthesis prevented Bax-precipitation and promoteed its insertion into nanodiscs. **(A)** BaxWT was synthesized alone or co-synthesized with Tom22 in the presence of 3.5µM nanodiscs (ND). The ratio of piVEX plasmids encoding the two proteins was 2 to 1 (typically 15ng and 7.5ng for Bax and Tom22, respectively). After the synthesis, the reaction mix was weighted and centrifuged at 20,000 x g for 15 minutes. The pellet was resuspended in the same volume of buffer B (25mM Hepes/Na pH 7.4, 200mM NaCl, 0.1mM EDTA). Aliquots from both the pellet (p) and supernatant (sn) were mixed with an equal volume of Laemmli buffer 2X, and analyzed by SDS-PAGE and western blot against Bax and Tom22. Note that Tom22 was visible under the forms of a monomer and a dimer. For the sake of clarity, only the monomer will be displayed in following experiments. **(B)** Quantitative analyzis of the effect of the co-synthesis of Tom22 on nanodisc-induced Bax precipitation. Non-saturated western-blots similar to **(A)** were quantified with Image J and the % of Bax in the pellets was calculated. Each point represents an independent synthesis assay. (****unpaired t-test, p<0.0001). **(C)** The remaining supernatant (typically 85µL) was mixed with 100µL of Ni-NTA-agarose (Qiagen) pre-washed in buffer B, and incubated for 2 hours at 4°C under gentle agitation. The mixture was centrifuged (2000 x g, 30 seconds) and the unbound fraction (lanes u) was collected. The resin was then washed 3 times with 500µL buffer W (20mM Tris/HCl pH 8.0, 500mM NaCl). An identical volume as the unbound fraction of buffer E (20mM Tris/HCl pH8.0, 200mM NaCl, 300mM imidazole) was added to recover Ni^2+^-bound nanodiscs) (lanes b). An identical volume of the unbound (u) and bound (b) fractions were mixed with an identical volume of Laemmli buffer 2X, and analyzed by SDS-PAGE and western-blot against Bax, Tom22 and His7-MSP1E3D1. **(D)** Quantification of Bax bound to nanodiscs (b). Non-saturated western-blots similar to **(C)** were quantified with Image J and the % of Bax in fraction b was calculated. Each point represents an independent synthesis assay. (**unpaired t-test, p<0.01). **(E)** Purified nanodiscs in the fraction “b” of **(C)** were dialyzed against buffer D (20mM Tris/HCl pH 8.0, 200mM NaCl) to remove imidazole. They were then added with 100mM Na _2_CO_3_, pH 10.0 and incubated for 15 minutes. Nanodiscs were then re-purified on Ni-NTA, as in **(C)**. **(F,G,H)** Same experiments as in **(A,C,E)** with Tom22(1-52) instead of full-length Tom22.

It has been reported that the mitochondrial receptor Tom22 was involved in Bax targeting to the MOM [15]. Strikingly, we observed that the co-synthesis of Tom22 with Bax prevented Bax precipitation (Fig.1A,B) and promoted the association of Bax to nanodiscs (Fig.1C,D).

To discriminate between loose protein/lipid interaction and membrane-inserted proteins, nanodiscs were submitted to an alkaline treatment to remove peripheral proteins. This treatment did not alter the stability of nanodiscs (Fig.S2A) and efficiently removed Kras, a known peripheral protein [37] (Fig.S2B). In the presence of Tom22, a large fraction of Bax remained associated to nanodiscs, showing that the co-synthesis with Tom22 promoted the membrane insertion of Bax (Fig.1E).

The alkaline treatment showed that Tom22 was itself inserted into nanodiscs (Fig.1E). We previously demonstrated that the co-synthesis of Bcl-xL, but not of Bcl-xLΔC, stimulated Bax insertion into nanodiscs [36]. We then asked if the insertion of Tom22 was also required for Bax insertion. Bax was co-synthesized with the cytosolic N-terminal domain of Tom22 (residues 1-52), that contains the negatively charged residues involved in the sorting of mitochondria-targeted proteins [38]. Like full-length Tom22, Tom22(1-52) prevented Bax precipitation (Fig.1F), and stimulated Bax association to nanodiscs (Fig.1D,G) and Bax insertion (Fig.1H). We concluded that, contrary to Bcl-xL, the membrane insertion of Tom22 was not required to promote Bax insertion. This showed that the stimulating effect of Tom22 on Bax insertion did not involve the co-insertion of the two proteins, but rather promoted a conformational change of Bax enabling its membrane insertion.

Importantly, the effects of Tom22 were specific: the co-synthesis of Bax with Tom20 did not prevent Bax precipitation nor promote Bax insertion (Fig.S2C,D), in agreement with the observation that the down-regulation of Tom20 did not change Bax mitochondrial localization and function *in cellulo* [15].

### Bax precipitation and Tom22 effects depend on the exposure of the hydrophobic GALLL motif in Bax Hα1

The interaction of soluble Bax with liposomes induces the exposure of the N-terminal epitope recognized by the 6A7 antibody [39], that is masked in soluble, inactive Bax [40]. Bax was then synthesized in the absence of nanodiscs, then added to nanodiscs: like liposomes, nanodiscs increased the exposure of the 6A7 epitope, to the same extent as the constitutively active mutant BaxR^9^E ([41]; see also below) in which the 6A7 epitope was constitutively exposed (Fig.2A).

**Figure 2.**
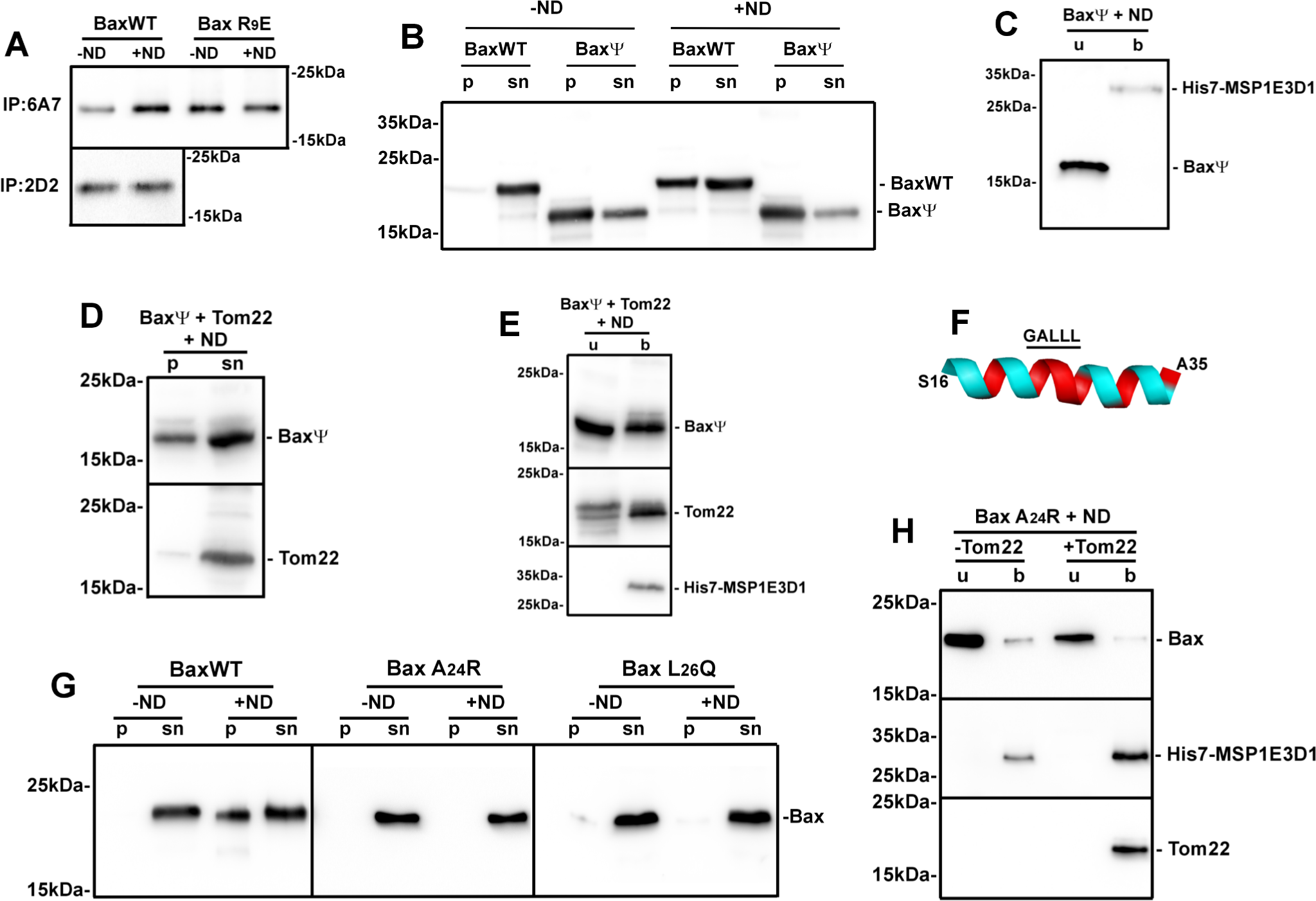
The GALLL motif in Hα1 was responsible for nanodisc-induced Bax precipitation, and interacted with Tom22. **(A)** BaxWT and the constitutively active mutant BaxR^9^E were synthesized in the absence of nanodiscs. Reaction mixes were centrifuged (20,000 x g, 15 minutes) and equal volumes of the supernatants were added or not with 3.5µM nanodiscs. 1µg anti-Bax 6A7 or 2D2 antibodies (Santa-Cruz Biotechnology) were added and samples were incubated overnight at 4°C under gentle agitation. 50µL Protein-G sepharose beads (Cytiva) were washed with buffer IP (20mM Tris/HCl pH 8.0, 200mM NaCl) and were added for 3 hours. The suspensions were then washed 5 times with 200µL of buffer WIP (25mM Tris/HCl pH 7.4, 500mM NaCl, 1mM EDTA, 1% Nonidet P-40, 10% glycerol) and twice with the same buffer diluted 10 times. Bound proteins were recovered by adding 30µL of Laemmli buffer (without β-mercaptoethanol) and heating at 65°C for 15 minutes. Samples were analyzed by SDS-PAGE and western-blot against Bax, with a distinct antibody from the ones used for immunoprecipitations (Abcam). The 2D2 antibody did not immunoprecipitate the Bax R^9^E mutant. **(B)** BaxWT and BaxΨ (deleted of residues 1-19) were synthesized in the absence or in the presence of nanodiscs and analyzed like in Fig.1A. Western-blots were done with the Abcam antibody of which the undisclosed epitope is away from the N-terminus. **(C)** Supernatants from BaxΨ in **(B)** were analyzed as in Fig.1C, showing the absence of BaxΨ-association to nanodiscs. **(D)** BaxΨ was co-synthesized with Tom22 in the presence of nanodiscs, and samples were analyzed like in **(B)**. **(E)** Supernatants from **(D)** were analyzed like in **(C)**, showing the partial association of BaxΨ to nanodiscs. **(F)** Representation of the Hα1 of Bax in the 1F16 structure [3]. Hydrophilic residues are in blue and hydrophobic residues are in red. **(G)** Same experiment as in Fig.1A with BaxWT, BaxA^24^R and BaxL^26^Q, showing the absence of precipitation of BaxA^24^R and BaxL^26^Q. **(H)** Same experiment as in Fig.1C with BaxA^24^R showing its absence of association to nanodiscs.

This antibody recognizes the epitope ^13^PTSSEQI^19^ of Bax only when it moves away from the core of the protein [42]. It is located within the ART sequence (Apoptotic Regulation of Targeting; residues 1-19), of which the full deletion stimulated Bax mitochondrial localization and its ability to release cytochrome c [12,14]. We then compared the cell-free synthesis of BaxΨ (carrying a deletion of residues 1-19) to full-length Bax (BaxWT). Contrary to BaxWT, BaxΨ precipitated in the absence of nanodiscs (Fig.2B), suggesting that ART deletion led to the exposure of a hydrophobic domain of the protein leading to its precipitation, mirroring the effect of nanodiscs on BaxWT. This was consistent with the hypothesis that the interaction of BaxWT with nanodiscs induced a movement of the ART sequence resulting in the exposure of hydrophobic domains leading to Bax precipitation. When synthesized in the presence of nanodiscs, the behavior of BaxΨ was the same as BaxWT: it precipitated (Fig.2B) and the small fraction remaining in the supernatant was not inserted (Fig.2C), unless co-synthesized with Tom22 (Fig.2D,E).

ART deletion exposes Hα1, and substitutions in this helix decrease the mitochondrial localization of both BaxΨ and BaxWT [14]. Hα1 contains a succession of hydrophilic and hydrophobic residues, with a longer motif of five hydrophobic residues ^23^GALLL^27^, thus displaying a hydrophobic face, and a hydrophilic face that is interrupted by a hydrophobic helix turn (Fig.2F). Since the exposure of the hydrophobic GALLL motif might be responsible for Bax precipitation, we replaced A^24^ or L^26^ by hydrophilic residues R and Q, respectively. Contrary to BaxWT, mutants BaxA^24^R and BaxL^26^Q did not precipitate when synthesized in the presence of nanodiscs (Fig.2G), showing that the hydrophobic GALLL motif was indeed involved in intermolecular interactions driving Bax molecules to precipitate following Hα1 exposure. Interestingly, when co-expressed with Tom22, BaxA^24^R was not inserted (Fig.2H) and BaxL^26^Q displayed an intermediate behavior between BaxWT and BaxA^24^R (Fig.S3). These data emphasized the role of the GALLL motif in Bax/Tom22 interaction, in agreement with the observation that Tom22 bound to GALLL-containing peptides [15].

### Bax activated by BH3 domains required Tom22 for membrane-insertion

*In cellulo*, the interaction between Bax and Tom22 only occurred after apoptosis had been triggered, suggesting that an initial conformational change of Bax was required [15]. BH3-only proteins Bid and Bim are the main activators of Bax. However, we observed that cBid, Bid-BH3 and Bim-BH3 peptides and the Bax-activator BTSA1 did not prevent nanodisc-induced Bax precipitation (Fig.S4A,B), nor stimulated Bax insertion into nanodiscs (Fig.3A,B). As expected, the co-synthesis of Tom22 prevented Bax precipitation (Fig.S4C) and restored Bax insertion in the presence of BH3 activators (Fig.S4D).

**Figure 3.**
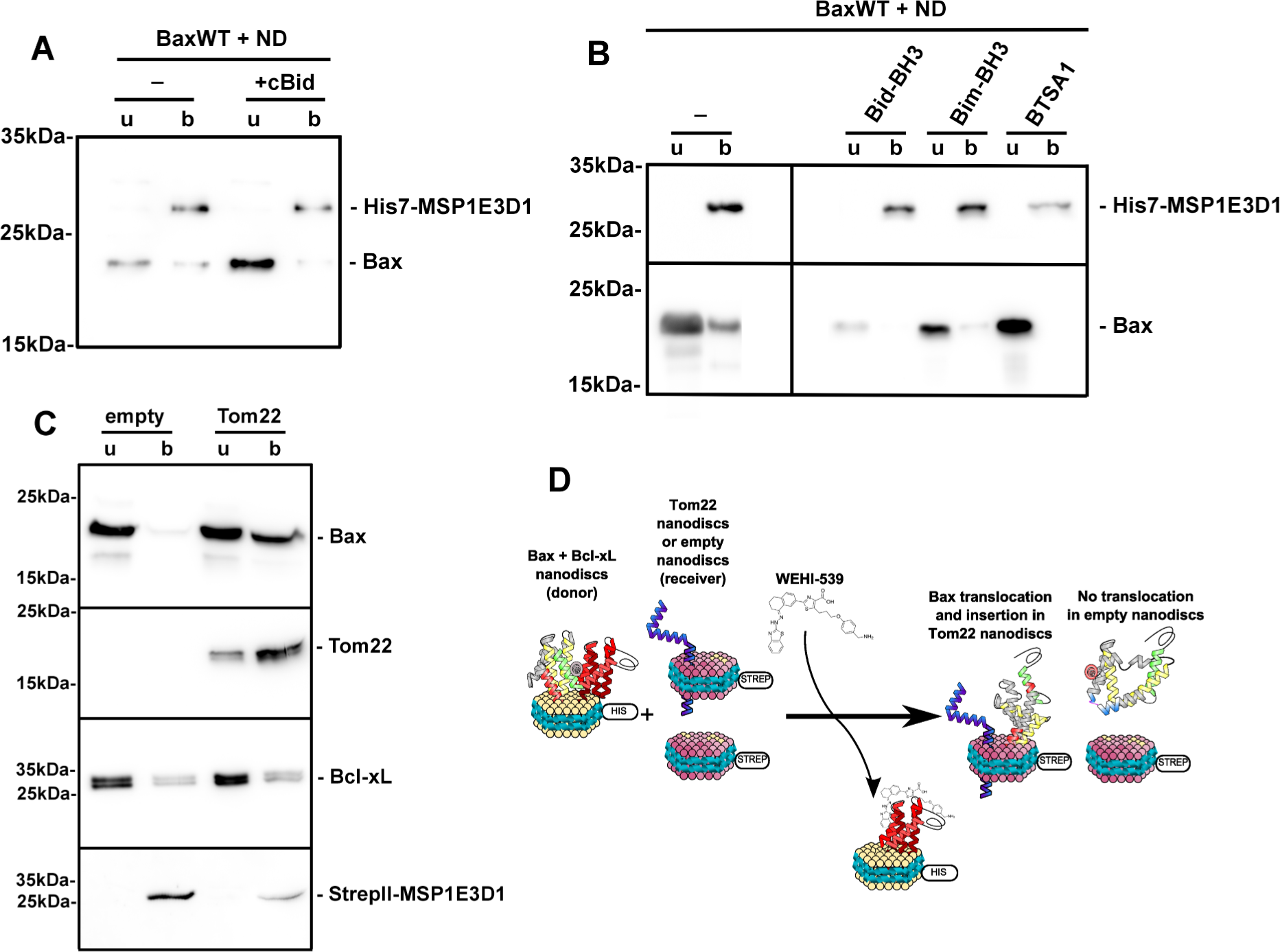
Tom22-dependent insertion of Bax activated by BH3-domains. **(A)** Same experiments as in Fig.1C (without Tom22), except that 10µg caspase-8-cleaved Bid (cBid) was included in the reaction mix. We determined that the maximal concentration of Bax produced in the cell-free system was 0.5 mg/mL, *i.e.* 50µg of Bax in the 100µL-reaction mix. We therefore set up a cBid to Bax ratio of ∼1 to 5 (considering that the sizes of the two proteins are close to each other) [67]. **(B)** Same experiment as in Fig.1C (without Tom22) in the presence of 1µM Bim-BH3 (PEIWIAQELRRIGDEFNAYYA), 1µM Bid-BH3 (ESQEDIIRNIARHLAQVGDSMDRSIPPG) (Genscript), or 1µM BTSA1 (Medchem) [74]. The concentration refers to the volume of the whole mix (reaction mix + feeding mix) because the sizes of the three molecules are below the cut-off of the dialysis membrane. **(C)** Bax and Bcl-xL were co-synthesized in the presence of nanodiscs made with His7-tagged MSP1E3D1, purified on Ni-NTA, and dialyzed against ND buffer (200 mM NaCl, 10mM Hepes/Na, pH 7.4) [36]. Tom22 was synthesized in the presence of nanodiscs made with StrepII-tagged MSP1E3D1, purified on Strep-Tactin (IBA), and dialyzed against ND buffer. Bax/Bcl-xL-containing His7-nanodiscs were mixed with the same amount of StrepII-nanodiscs, either empty or containing Tom22. 250nM of WEHI-539 were added and the mixture was incubated overnight at 4°C. StrepII-nanodiscs were purified again on Strep-Tactin and the unbound and bound fractions were analyzed. **(D)** Schematic representation of the results of **(C)**.

In cancer cells, the overexpression of anti-apoptotic proteins, such as Bcl-xL, prevents Bax activation. This inhibition is relieved by BH3-only proteins, and by BH3-mimetic compounds ([43] for review), releasing Bax under a primed conformation [44]. We set up a nanodisc-to-nanodisc transfer experiment to appraise if Tom22 was still required when Bax was primed by its interaction with Bcl-xL then released by the BH3-mimetic WEHI-539. Bax and Bcl-xL were co-synthesized and co-inserted in nanodiscs formed with His7-tagged MSP1E3D1 [36]. These nanodiscs were purified and incubated with nanodiscs formed with StrepII-tagged MSP1E3D1, either empty or containing Tom22. The addition of WEHI-539 was expected to break the interaction between Bax and Bcl-xL, therefore releasing Bax under a primed conformation. The presence of Bax was then measured in purified StrepII-tagged nanodiscs (Fig.3C). We observed that Bax was translocated to StrepII-nanodiscs when Tom22 was present, but not to empty StrepII-nanodiscs. This showed that the presence of Tom22 was required for Bax insertion after it was primed through its interaction with Bcl-xL followed by its release by WEHI-539 (Fig.3D).

These data showed that, even after Bax had been activated by its interaction with its partners, either directly by Bim or Bid, or indirectly by its interaction with Bcl-xL then release by WEHI-539, the interaction with Tom22 was still required for its insertion into nanodiscs.

### Tom22 triggers a conformational change leading to the extrusion of Bax Hα9

Our data suggested that a two-step process initiated Bax insertion into nanodiscs: (i) the interaction of Bax with the lipid bilayer induced a movement of ART exposing GALLL motif in Hα1 and (ii) Tom22 interacted with the exposed GALLL motif, thus triggering conformational changes leading to Bax membrane insertion. We next investigated the nature of these conformational changes.

Because the stability of the soluble conformation of Bax depends on the embedding of its C-terminal hydrophobic Hα9 in the hydrophobic groove, Bax insertion first requires the extrusion of Hα9 from the groove [3]. To investigate the effect of Tom22 on Hα9 mobility, we measured the exposure of a single cysteine residue introduced at position 177. Like BaxWT, the triple mutant BaxC ^62^S/C^126^S/V^177^C was dependent on Tom22 for its insertion into nanodiscs (Fig.4A). We next measured the accessibility of the V^177^C residue to NEM-PEG in the different conformations stabilized by our assay (Fig.4B). In the absence of nanodiscs, V^177^C was accessible to NEM-PEG. When synthesized in the presence of nanodiscs, V^177^C residue of the soluble fraction of Bax remained exposed, regardless of the co-synthesis of Tom22. On the contrary, V^177^C of nanodisc-inserted Bax (in the presence of Tom22) was no longer accessible, showing that Hα9 was inserted into the lipid bilayer.

**Figure 4.**
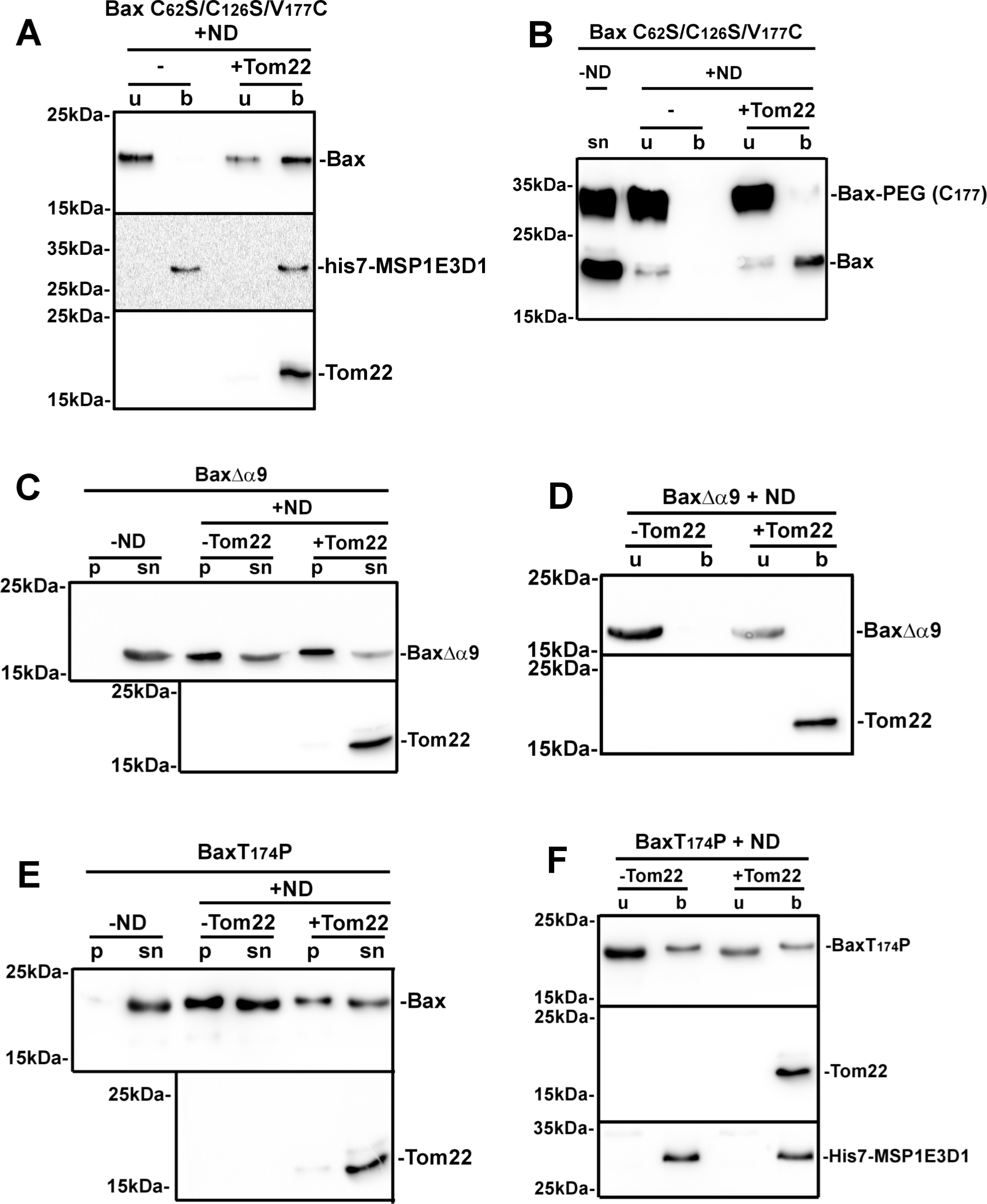
Effect of nanodiscs and Tom22 on the exposure and insertion of Hα9. **(A)** Same experiment as in Fig.1C with the mutant BaxC^62^S/C^126^S/V^177^C, in which the two endogenous cysteine residues in positions 62 and 126 have been replaced by serine, and a cysteine has been introduced in Hα9. **(B)** Fractions from **(A)** were incubated for 1 minute in the presence of 0.2mM NEM-PEG (5kDa) and the reaction was stopped by adding 20mM NEM and a 10-minutes additional incubation. The lane “-ND” is a synthesis without ND and Tom22. Samples were then analyzed by SDS-PAGE and western-blot against Bax. **(C,E)** Same experiments as in Fig.1A on mutants BaxΔα9 and BaxT^174^P, showing that Tom22 did not prevent their precipitation. **(D,F)** Same experiments as in Fig.1C on mutants BaxΔα9 and BaxT^174^P showing that Tom22 did not promote their insertion.

We next showed that the insertion of Hα9 was a prerequisite for Tom22-induced membrane-insertion of Bax, and the protection against nanodisc-induced precipitation. For this, we analyzed Bax deprived of Hα9 (BaxΔα9, residues 1-169). Like full-length Bax, and contrary to BaxΨ, BaxΔα9 was soluble in the absence of nanodiscs and precipitated in their presence (Fig.4C). However, its precipitation was not prevented by the co-synthesis with Tom22 (Fig.4C). As expected, BaxΔα9 was not inserted in nanodiscs (Fig.4D). These data confirmed that the rescue of Bax precipitation by Tom22 was actually related to the Hα9-dependent insertion of Bax.

According to the Cosmic database (https://cancer.sanger.ac.uk/cosmic), a T^174^P substitution in Hα9 is frequently found in squamous cell carcinomas of the upper digestive track [45]. We constructed a mutant carrying this substitution, thus keeping Hα9 length, but breaking its helical conformation. The co-synthesis of Tom22 did not significantly rescue the precipitation of BaxT^174^P (Fig.4E) and did not stimulate its poor membrane-insertion (Fig.4F). This discarded the possibility that a side effect linked to the full deletion of Hα9 was responsible for nanodisc-induced BaxΔα9 precipitation, and showed that defective Bax insertion might be involved in defective apoptosis in cancer cells containing the T^174^P substitution.

These data established that the effects of Tom22, both on the prevention of Bax precipitation and the stimulation of Bax insertion into nanodiscs, strictly depended on the presence of a Hα9 able to insert into the bilayer. This was consistent with a model where, following the initial interaction of Bax with the lipid bilayer exposing the GALLL motif in Hα1 (without any conformational change at the C-terminus at this stage), the subsequent interaction of Tom22 with the GALLL motif promoted the extrusion of Hα9 followed by its insertion. This later step could not occur in BaxΔα9 and BaxT^174^P mutants, resulting in their precipitation caused by the hydrophobic interaction between GALLL motifs.

### Involvement of Bax R^9^ and D^154^ in Tom22-induced Hα9-insertion

We have previously reported that soluble, inactive Bax might be stabilized by an electrostatic interaction between residues R^9^ in the ART and D^154^ in Hα7: indeed, single charge inversions of either one of these residues (R^9^E or D^154^K) resulted in their constitutive mitochondrial localization and permeabilization of the MOM to cytochrome c, while the double-substituted mutant R ^9^E/D^154^K, in which the electrostatic interaction was restored, displayed the same soluble, inactive conformation as BaxWT [41]. We therefore tested the active mutant R^9^E and the inactive revertant R^9^E/D^154^K in our set-up.

Contrary to BaxWT, BaxR^9^E did not precipitate when synthesized in the presence of nanodiscs (Fig.5A) and, strikingly, was spontaneously associated to nanodiscs (Fig.5B). As expected, this was reversed by the introduction of the D^154^K mutation (Fig.5A,B). Next, the R^9^E substitution was introduced in the triple mutant BaxC^62^S/C^126^S/V^177^C described above. NEM-PEG labeling showed that Hα9 was actually inserted in the lipid bilayer (Fig.5C). A BaxR^9^E/Δα9 mutant showed that the deletion of Hα9 impaired both the protective effect of the R^9^E mutation on Bax precipitation (Fig.5D) and its association to nanodiscs (Fig.5E). These data showed that the BaxR^9^E mutant synthesized alone displayed the same behavior as BaxWT co-synthesized with Tom22, and thus fully recapitulated the conformational changes initiated by the interaction of Bax with Tom22, leading to the insertion of Hα9. The presence of a D^154^Y substitution in some tumors has been reported in Uniprot (https://www.uniprot.org/uniprotkb/Q07812/variant-viewer) and in the Spanish database Intogen (https://www.intogen.org/search?gene=BAX). Like for BaxWT, the co-synthesis of Tom22 protected BaxD^154^Y against nanodisc-induced precipitation (Fig.5F). However, contrary to BaxWT, Tom22 did not promote BaxD^154^Y insertion (Fig.5G). From these data, we concluded that the substitution D^154^Y did not impair the interaction of Bax with Tom22 and the associated rescue of Bax precipitation, but rather prevented Tom22-induced conformational change leading to Bax insertion. Like for the mutant T^174^P described above, this behavior might relate to a defective apoptotic process in cancer cells displaying the D^154^Y substitution.

**Figure 5:**
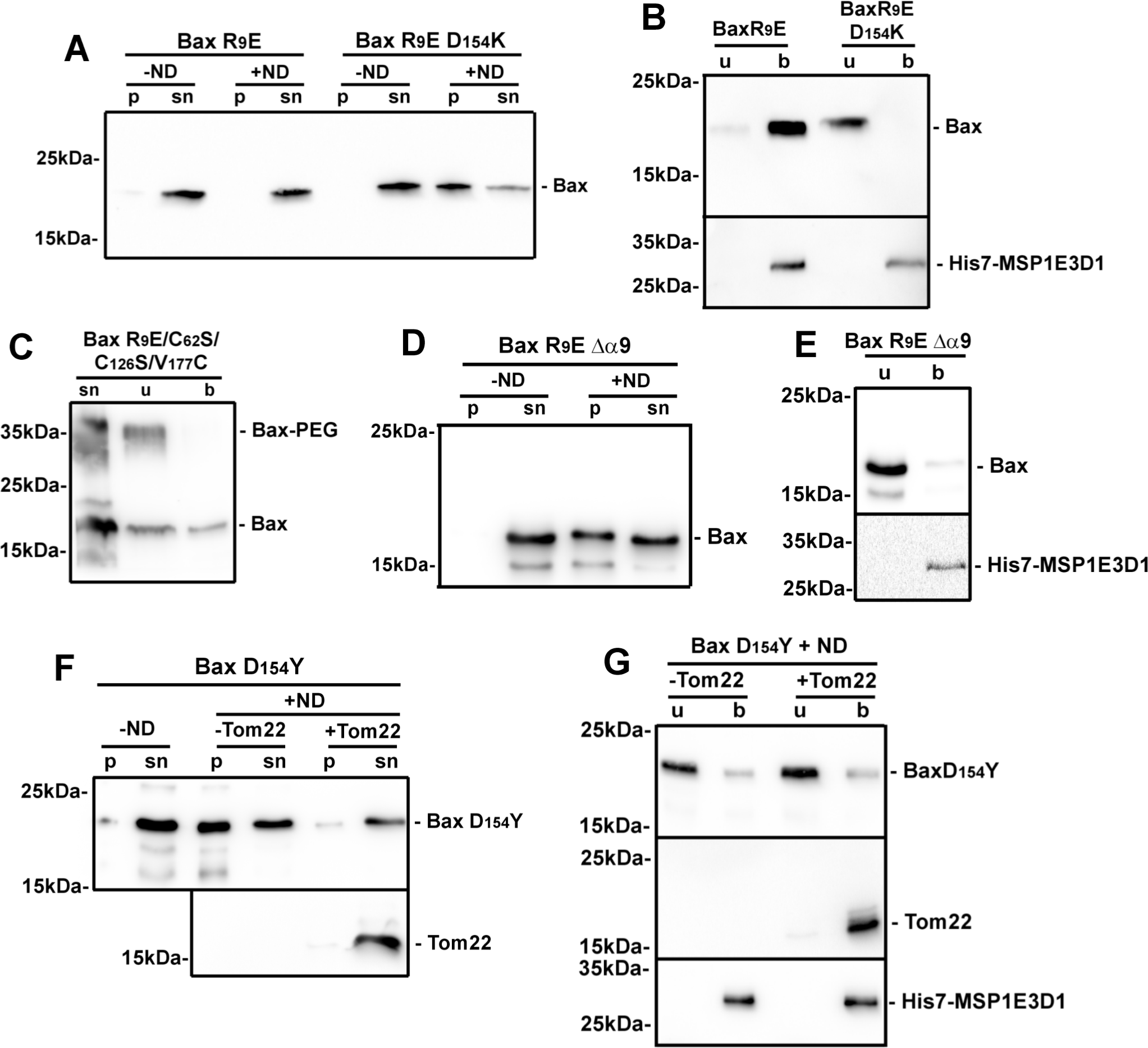
Effects of substutions on R^9^ and D^154^ residues. **(A,B)** Same experiments as in Fig.1A and 1C on mutants BaxR^9^E and BaxR^9^E/D^154^K in the absence of Tom22, showing that BaxR^9^E was spontaneously inserted, and that the mutation D^154^K reversed the spontaneous insertion. **(C)** Same experiment as in Fig.3B on mutant BaxR^9^E/C^62^S/C^126^S/V^177^C synthesized in the absence of Tom22, showing the spontaneous insertion of Hα9. **(D,E)** The mutant BaxR^9^E/Δα9 was synthesized in the absence of Tom22 and analyzed as in Fig.1A and 1C, showing that the deletion of Hα9 impaired the absence of precipitation and prevented the insertion induced by the mutation R^9^E. **(F,G)** Same experiments as in Fig.1A and 1C on mutant BaxD^154^Y, showing that Tom22 prevented its precipitation but did not promote its insertion.

### Pore-forming activity of Bax inserted through Tom22-mediated conformational change

Cell-free synthesis was done in the presence of liposomes loaded with Dextran-FITC (10kDa). An anti-FITC antibody was added to quench the fluorescence of released Dextran-FITC, and the fluorescence decay was followed during the synthesis of Bax, alone or co-synthesized with Tom22(1-52). The co-synthesis with Tom22(1-52) increased the rate of fluorescence decay, compared to BaxWT alone (Fig.6A,B). This suggested that Tom22(1-52) not only stimulated the insertion of BaxWT but further stimulated its ability to form pores permeable to a molecule having a similar size as cytochrome c (12.5kDa). BaxR^9^E synthesized alone without Tom22 also exhibited a higher rate of permeabilization, consistent with the above experiments showing that the R^9^E substitution recapitulated the effect of Tom22. These experiments showed that both BaxWT co-synthesized with Tom22(1-52) and BaxR^9^E synthesized alone formed pores large enough to release cytochrome c.

**Figure 6.**
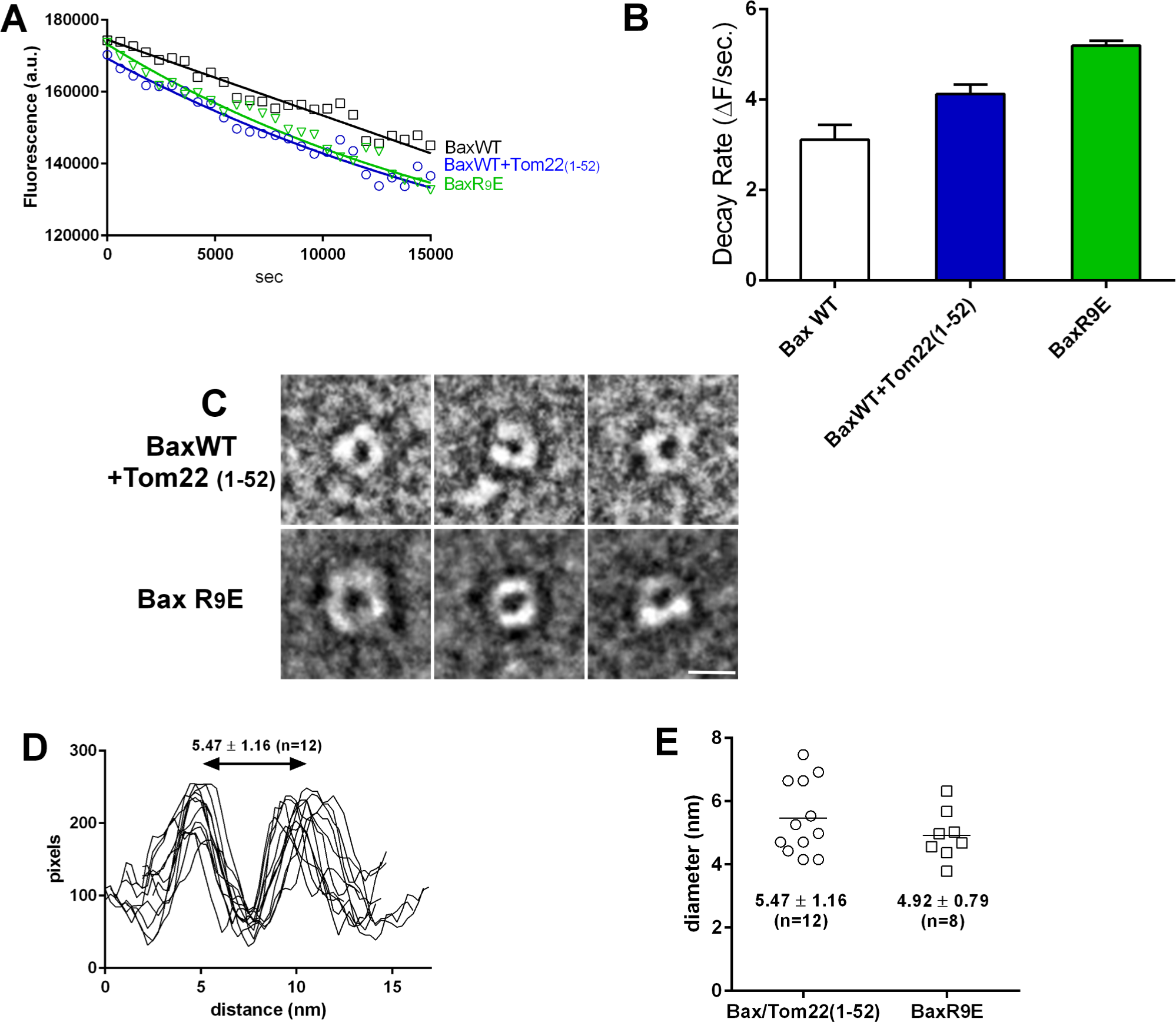
Effect of Tom22 and of the mutation BaxR^9^E on Bax pore forming capacity. **(A)** BaxWT was synthesized alone or co-synthesized with Tom22(1-52), and the mutant BaxR^9^E was synthesized alone. 20µL liposomes loaded with Dextran-FITC (10kDa) (Sigma) and 4µg of a fluorescence quenching anti-FITC antibody (ThermoFisher Scientific) were added in the reaction mix. Mixes were put in a 96-well fluorescence plate, and incubated at 28°C in a thermostated fluorescence plate reader (Clariostar, BMG Labtech). Samples fluorescence was recorded at 10-minutes intervals (ex: 475-490 nm; em: 515-545 nm). **(B)** Inital rates of fluorescence decays in experiments of **(A)**. Plots were fitted with exponential decay and the rate constants were calculated and averaged from two independent experiments. R^2^ values were 0.95, 0.94 and 0.98 for BaxWT, BaxWT+Tom22(1-52) and BaxR^9^E, respectively. **(C)** BaxWT was co-synthesized with Tom22(1-52) and BaxR^9^E was synthesized alone in the presence of nanodiscs. Nanodiscs were purified and concentrated on Vivaspin concentrators (cut-off 50kDa; Sartorius). They were negatively stained with uranyle acetate and observed by TEM (see methods). Selected images showing the presence of a pore are shown. The scale bar is 10nm. **(D,E)** Pixel densities were measured on images along a line crossing the largest diameter of the pores.

TEM of negatively stained nanodiscs showed that both Bax/Tom22(1-52) and BaxR^9^E could form a structure resembling a pore (Fig.6C) with a diameter of about 5 nm (Fig.6D,E). This size was consistent with both electrophysiology experiments on MOM [46,47] and reconstituted oligomers [46], and with TEM observations of precipitated/resolubilized oligomers of BaxP^168^A reconstituted in nanodiscs by the co-formation method [32]. These data showed that the direct nanodisc insertion of BaxWT co-synthesized with Tom22, as well as that of BaxR^9^E synthesized alone, accurately embodied a pore suitable for the release of cytochrome c.

## Discussion

### Tom22-induced conformational changes lead to Bax membrane-insertion

Setting up a model combining the cell-free synthesis of Bax to its insertion into nanodiscs led us to reappraise the involvement of the mitochondrial receptor Tom22 in Bax membrane insertion. We had previously reported that the presence of nanodiscs unexpectedly induced the precipitation of Bax [32]. Here, we report that the co-synthesis of Tom22 prevented this precipitation and promoted the insertion of Bax into nanodiscs (Fig.1). To decipher the mechanisms underlying the effect of Tom22, we first investigated the molecular events leading to nanodisc-induced Bax precipitation.

The interaction of Bax with nanodiscs induced the exposure of the GALLL motif in Hα1 of Bax molecules resulting in their precipitation through hydrophobic interactions: indeed, substitutions to more hydrophilic residues (A^24^R, L^26^Q) prevented Bax precipitation (Fig.2G). Conversely, the deletion of ART, that constitutively exposes Hα1 [14], induced the precipitation of Bax, even in the absence of nanodiscs (Fig.2B).

The same GALLL motif is involved in the MOM targeting of Bax [14] and in the interaction between Bax and Tom22 [15]. Knocking down Tom22, or inactivating it by proteolysis or by a specific antibody, decreased Bax mitochondrial localization and apoptosis [15]. In the same study, peptide-mapping identified ^21^KTGALLLQ^28^ as the main domain of interaction of Bax with Tom22. However, the involvement of Tom22 in Bax mitochondrial targeting has been debated, and contradictory conclusions have been drawn [15–20]. The present report, based on a minimal model containing only Bax, Tom22 and a lipid bilayer stabilized in nanodiscs, unambiguously demonstrated that Tom22 not only prevented Bax precipitation but further promoted its insertion into nanodiscs (Fig.1C-E). As summarized in Fig.7A, once the GALLL motif was exposed, it could interact with Tom22 which triggered Bax membrane insertion. In the absence of Tom22, the hydrophobic interaction between GALLL motifs led to Bax precipitation. Fig.7B illustrates the consequence of mutations A^24^R and L^26^Q, which prevented Bax precipitation by decreasing the hydrophobicity of the GALLL motif but impaired the interaction with Tom22, while the deletion of ART constitutively exposed the GALLL motif leading to BaxΨ precipitation regardless of the presence of nanodiscs, but to its insertion into nanodiscs when Tom22 was present.

**Figure 7.**
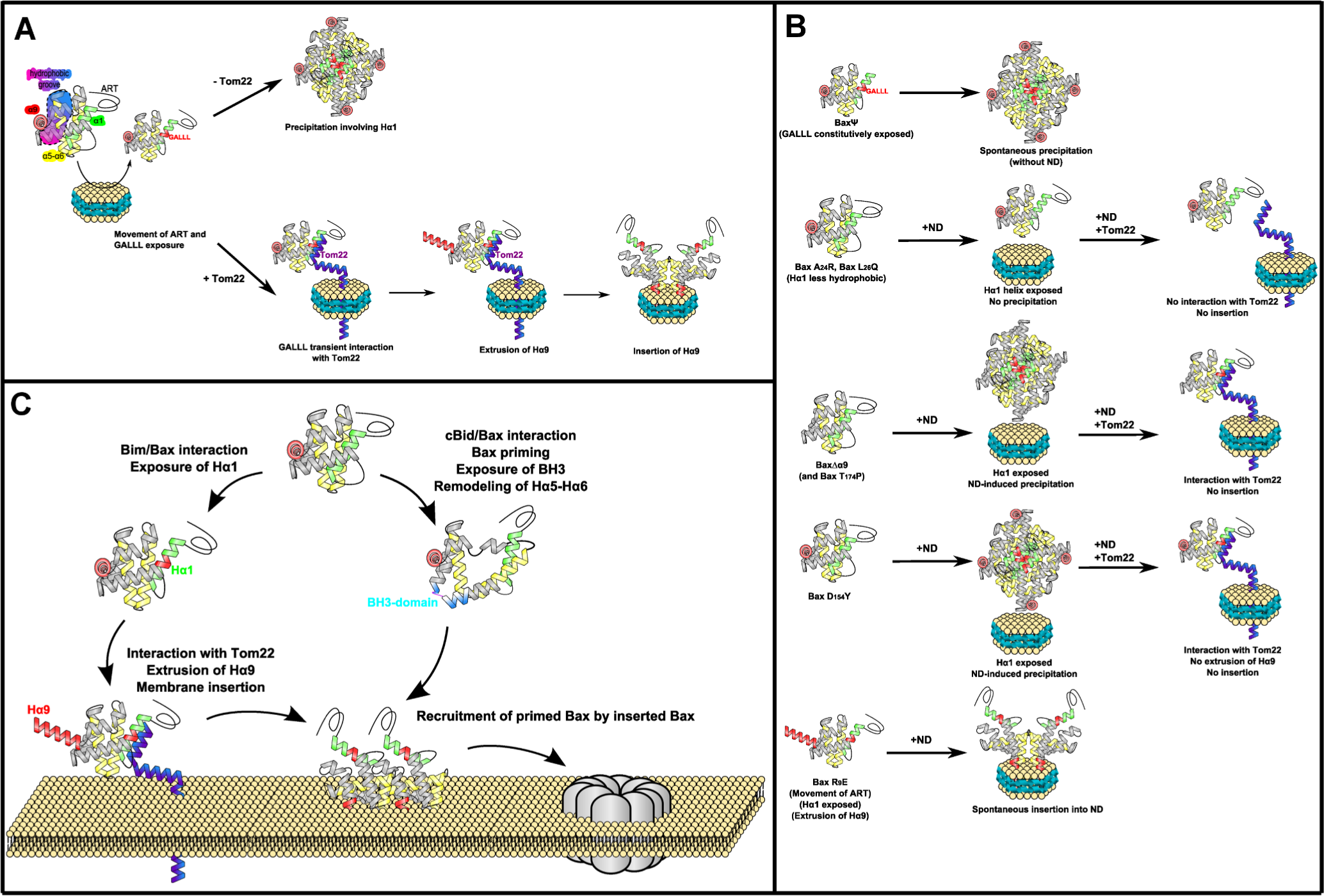
Representation of Bax interaction with nanodiscs and Tom22. **(A)** The interaction of Bax with nanodiscs induced the exposure of the GALLL motif in Hα1. In the absence of Tom22, interaction between GALLL motifs of different Bax molecules led to their precipitation. In the presence of Tom22, GALLL motif interacted with Tom22, promoting a conformationnal change leading to the extrusion, followed by the insertion of Hα9. **(B)** Behavior of the different mutants investigated in this study. - In BaxΨ, Hα1 was constitutively exposed, leading to its precipitation in the absence of nanodiscs, caused by hydrophobic interactions between GALLL motifs. - In BaxA^24^R and BaxL^26^Q, the exposure of Hα1 did not induce Bax precipitation in the presence of nanodics; however these mutants failed to interact with Tom22 and therefore were not inserted. - Like BaxWT, the mutants BaxΔα9 and BaxT^174^P exposed the GALLL motif in the presence of nanodiscs, and interacted with Tom22. However, in the absence of an adequate Hα9, they were not correctly inserted into nanodiscs - The mutant D^154^Y, like BaxWT, was precipitated in the presence of nanodiscs, and could interact with Tom22; however, it did not support the conformational change leading to Bax insertion. - The interaction of mutant R^9^E with nanodiscs led to the conformational change leading to the extrusion and insertion of Hα9 even in the absence of Tom22. **(C)** Hypothetic model of the concerted involvement of BH3-only proteins and Tom22 in Bax insertion in the MOM. The interaction of Bim with Bax at the non-canonical binding site promoted the exposure of Hα1, allowing the interaction of Bax with Tom22. Bax/Tom22 interaction led to the extrusion and insertion of Hα9. The interaction of cBid with Bax induced the exposure of the BH3-domain and the remodeling of the Hα5/Hα6 [69,70], allowing the recruitment of cBid-activated Bax by membrane-inserted Bax, leading to the formation of membrane-inserted oligomers.

The membrane insertion of Bax requires the extrusion of Hα9 from the hydrophobic groove ([3,4], [48] for review). It has been proposed that the interaction of Bim at the non-canonical binding site, including Hα1, Hα6 and the α1-α2 loop [8] could trigger long range conformational changes leading to Hα9 extrusion [9], which was further supported by molecular dynamics simulations [49] and by the observation that mutations in the α1-α2 loop prevented the extrusion of Hα9 [10]. We observed that the interaction of Tom22 with exposed Hα1 led to the insertion of Hα9 (Fig.4B). Conversely, the deletion of Hα9 or the introduction of the helix-breaking mutation T^174^P abolished the protective effect of Tom22 on Bax precipitation (Fig.4C,E). This showed that, after Hα1 had been exposed, any failure of Hα9 membrane-insertion resulted in Bax precipitation, further emphasizing the crucial role of Hα9-extrusion as the key step towards Bax insertion.

Although Bax is not generally considered as a mutational hotspot in cancers, several misense mutations found in tumors might alter its pro-apoptotic function [50,51]. Furthermore, substitutions that had been primarily generated to study Bax conformational changes targeted residues later found to be mutated or modified in some cancers. Namely, we have reported the possible occurrence of an electrostatic interaction between R^9^ and D^154^ stabilizing soluble Bax [41]. Interestingly, BaxR^9^ can be monomethylated [52], which is expected to destabilize the interaction with D^154^. A D^154^Y substitution has also been found in some tumours (https://www.uniprot.org/uniprotkb/Q07812/variant-viewer; https://www.intogen.org/search?gene=BAX). This led us to investigate the consequences of substitutions of these two interacting residues. The R^9^E mutant did not precipitate in the presence of nanodiscs (Fig.5A) and was spontaneously inserted (Fig.5B,C) in a way that depended on Hα9 (Fig.5D,E). This mutation fully recapitulated the effect of Tom22, inducing Bax spontaneous insertion, regardless of the presence of Tom22 (Fig.7B).

The D^154^Y substitution did not prevent the rescue of Bax precipitation by Tom22 (Fig.5F), but impaired Bax insertion (Fig.5G) (Fig.7B). This suggested that the Tom22-triggered conformational change leading to Hα9 insertion involved D^154^. Docking simulations between Bax and Tom22 have suggested that Tom22 might interact more strongly with soluble Bax than with Hα9-inserted Bax [53]. In the same study, simulating the breaking of the hydrogen bond between S^184^ and D^98^ resulted in the movement of the C-terminus of Bax, starting at D^154^. These data were consistent with the hypothesis that a transient interaction between Tom22 and soluble Bax was followed by a movement of the C-terminus of Bax around D^154^, leading to Hα9 insertion and then the release of membrane-inserted Bax from the interaction with Tom22.

### Tom22-dependent Bax pore formation

Both BaxWT co-synthesized with Tom22 and BaxR^9^E synthesized alone permeabilized liposomes to dextran-10kDa (Fig.6A,B) and formed nanodiscs containing a pore displaying a ∼5nm diameter (Fig.6C-E), which was consistent with electrophysiology data on MOM [46,47,52] and reconstituted Bax oligomers [46]. This size was much smaller than the one of large structures observed by high resolution fluorescence microscopy and atomic force microscopy, reaching 24 to 176 nm in diameter [55–57], which are compatible with the formation of megapores involved in the release of mtDNA during inflammatory responses [58,59]. Conversely, the smaller 5nm-pore might correspond to early apoptotic steps, when small apoptogenic factors, such as cytochrome c (∼3nm) and smac/diablo (∼3.5nm) are released. It was however somewhat larger than the 3.5nm size that had been measured in nanodiscs containing Bid-BH3/low pH-activated Bax [30]. On the basis of cryoEM images and molecular dynamics simulations, it was proposed that Bax monomers (or possibly dimers) might form lipidic pores by rearranging phospholipid molecules in nanodiscs [31]. NMR structural studies of the Bax core (Bax Hα2-Hα5) in bicelles are compatible with a pore of which the walls were made of Bax dimers interacting with lipids, which might be further organized in dodecamers to generate a pore large enough to promote the release of cytochrome c [60]. Electrophysiology data showed that Bax pores exist under different forms displaying different conductance substates [61], which might accurately describe the continuum of Bax-induced permeabilization to small apoptogenic factors such as cytochrome c and smac/diablo, then to larger apoptogenic factors AIF and endonuclease G, and up to massive permeability events leading to the release of mitochondrial DNA in inflammatory responses ([62–64] for reviews). However, the conformation of membrane Bax in “small apoptotic pores” and “large inflammation pores” might not be identical. Although they exhibit a certain degree of flexibility [65], the nature of nanodiscs imposes more physical constraints, both in terms of size and lipid packing, than, for example, liposomes [66]. Bim or cBid are sufficient to insert Bax into liposomes [35,67], but not into nanodiscs (Fig.3A,B). It is tempting to speculate that this might reflect different types of pores formed from Bax monomers/dimers having distinct conformations and, owing to the physical constraints they impose, nanodiscs may provide a mean to stabilize small pores *vs* megapores.

### On the respective roles of BH3 direct activators and Tom22 on Bax membrane insertion and pore formation

The induction of apoptosis is a prerequisite for Bax/Tom22 interaction *in cellulo* [15]. It can therefore be hypothesized that BH3 activators and Tom22 might cooperate during the process of Bax activation. The concerted action of Bim and Tom22 would lead to the extrusion then membrane insertion of Bax (Fig.7C). Once inserted, Bax could actually play the role of a receptor to itself [68]. The interaction of Bax with cBid, that induces the exposure of the BH3-domain and the conformational change of Hα5/Hα6 [69], might therefore prime soluble Bax for its interaction with membrane-inserted Bax, leading to the step-wise formation of oligomers [70] (Fig.7C).

In our minimal model, BH3 activators were not required, and the interaction of Bax with the lipid bilayer seemed sufficient to expose Hα1. In cancer cells, Bax activation by BH3 proteins is limited by the overexpression of anti-apoptotic proteins, such as Bcl-xL, leading us to investigate if Tom22 was still involved in Bax activation in the presence of Bcl-xL. Following the rupture of Bax/Bcl-xL interaction by WEHI-539, Bax could be translocated from Bcl-xL-containing nanodiscs and inserted into new nanodiscs only when Tom22 was present (Fig.3C,D). This demonstrated that Tom22 was required for Bax membrane insertion after it was primed by the engagement then release of its interaction with Bcl-xL [44]. This emphasized the crucial role of Tom22 in Bax membrane-insertion under conditions reflecting the context of cancer cells. This might be fully relevant in the context of the use of BH3-mimetics as anti-tumor agents.

### Conclusion

Since the initial report showing that Tom22 was involved in the MOM-targeting of Bax [15], no mechanistic explanation had been provided and the actual role of Tom22 had been questioned [19,20]. The present study connects the interaction of Bax with Tom22 to the movement of Hα9, as the primary event leading to Bax membrane insertion. It also explains why Bax could not be spontaneously inserted in nanodiscs [32]. This study paves the path for future structural investigations on Bax in nanodiscs, in which Bax forms a relatively small pore, consistent with the limited release of small proteins such as cytochrome c. We also propose a model combining the action of Tom22 and BH3-activators Bim and cBid on Bax insertion, and extend our observations to the role of Tom22 in the membrane-insertion of Bax after it is released from its inhibition by its anti-apoptotic partner Bcl-xL.

## Materials and methods

### Plasmids construction and cell-free protein synthesis

Full-length, untagged Bax, Tom22, Tom20 and Bcl-xL were cloned in the Nde1/Xho1 sites of the pIVEX 2.3a plasmid. Kras4b was cloned in the pIVEX 2.4d plasmid in frame with the N-terminal His6 tag. A double TAA stop codon was included to secure translation arrest during *in vitro* synthesis. Site-directed mutagenesis was done by an optimized Quickchange protocol [71] and mutations were verified by sequencing (Eurofins).

The plasmid pET28b encoding His7-MSP1E3D1 (Addgene) was transformed into *E. coli* BL21DE3 and the protein was purified as described previously [32,36]. A strepII-tagged version of the protein has been constructed by site-directed mutagenesis. The pAR1219 encoding T7-RNA polymerase was transformed into E.coli BL21DE3*, and the protein was purified according to [72]. Nanodiscs were formed with a mixture of phospholipids (48% POPC, 32% POPE, 12% DOPS, 8% cardiolipid (w/w)) (Anatrace). The cell-free synthesis protocol was detailed previously [32,36,72]. Syntheses were done in inverted microtubes containing 100µL of Reaction Mix in the round compartment of the cap and 1700µL of Feeding Mix, separated by a dialysis membrane (cut-off 8-10kDa), overnight at 28°C under gentle agitation (80 rpm).

After the syntheses, the volume of the reaction mixes was carefully measured and, after a 20,000 x g centrifugation, the pellet was re-suspended in the same volume, so that the same proportions of pellets and supernatants were loaded on SDS-PAGE. After the separation of nanodiscs and soluble proteins on Ni-NTA (Qiagen), the same proportions of unbound material and imidazole-eluted material were analyzed (Fig.S5). It follows that, for each gel, the amount of proteins in pellets (p) *v/s* supernatants (sn), and in unbound fraction (u) *v/s* bound fraction (b) are directly comparable.

### SDS-PAGE and Western blots

Proteins were solubilized in Laemmli buffer 2X and heated at 65°C for 20 minutes. Samples were loaded on SDS-PAGE (Mini Protean, Biorad). Tris-glycine gels were made of 12.5% of acrylamide/bisacrylamide (40/1). Gels were run at 15mA per gel. Protein transfer on PVDF membranes (Hybond, Cytiva) was done with the Transblot system (Biorad) for 1h30 at 350mA. Western-blotting was done in PBST or TBST, depending on the antibody (Table S6). Western-Blots were revealed by chimioluminescence (Luminata Forte, Millipore) and recorded with a digital camera (G-Box, Syngene).

### NEM-PEG labeling

NEM-PEG labeling was done according to [10]. Protein samples were incubated with 0.2mM methoxypolyethyleneglycol maleimide (NEM-PEG, 5kDa; Sigma-Aldrich) for 1 minute on ice. The reaction was stopped by adding 20mM NEM for 10 minutes.

### Dextran-FITC release from liposomes

Liposomes permeabilization was measured through the quenching of external Dextran-FITC by an anti-FITC antibody. A cell-free synthesis reaction was set-up in 100µL, in the presence of 10µL liposomes loaded with Dextran-FITC (10kDa, Sigma) (liposomes preparation is detailed in [32]. 4µg of anti-FITC antibody (Rabbit Monoclonal 6HC5LC9; ThermoFisher Scientific) was added to quench external FITC fluorescence. The mix was then added to black 96-well plates and incubated at 28°C in a fluorescence plate reader (Clariostar, BMG Labtech). Fluorescence was measured at 10-minutes intervals (ex: 475-490 nm, em: 515-545 nm).

### Bax translocation between nanodiscs

Bax and Bcl-xL were co-synthesized in the presence of nanodiscs formed with His7-tagged MSP and purified on Ni-NTA and dialyzed in ND buffer (200mM NaCl, 10mM Hepes, pH 7.4) to eliminate imidazole [36]. In parallel experiments, Tom22 was synthesized in the presence of nanodiscs formed with StrepII-tagged MSP, purified on Strep-Tactin (IBA) and dialyzed in ND buffer to eliminate biotin. The eluate from Ni-NTA containing Bax/Bcl-xL in His7-nanodiscs was mixed with an equal amount of StrepII-nanodiscs, empty or containing Tom22 (1.13 µM of each type of nanodiscs, expressed as MSP concentration). 250nM WEHI-539 (MedChem) was added to the mixes, that were incubated overnight at 4°C. Mixes were then loaded on Strep-Tactin and the unbound and bound fractions were collected and analyzed by western-blot.

### Dynamic Light Scattering

Dynamic Light Scattering (DLS) was measured in a DynaPro Nanostar (Wyatt) on 10µL samples. Triplicates of 10-acquisition measurements were done and averaged for each point.

### Transmission Electron Microscopy

Transmission Electron Microscopy (TEM) was done as described previously (Rouchidane Eyitayo et al., 2023a). Nanodiscs were concentrated on Vivaspin concentrators (50kDa, Sartorius). Negative staining was done according to [73]. Samples were observed on a Tecnai F20 FEG electron microscope (FEI, ThermoFisher Scientific) operated at 200 kV using an Eagle 4k_4k camera (ThermoFisher Scientific). Measurements were manually done with Image J (https://imagej.net/ij/) on dm3 files.

### Miscellaneous

Unless indicated otherwise, chemicals were from Clinisciences and Sigma-Aldrich.

Displayed blots are representative of at least 3 independent syntheses. Where indicated, quantification has been done on non-saturated blots and statistical analyses were done with GraphPad 6.05 tools.

## Acknowledgments

This work was supported by the CNRS, the University of Bordeaux and the Ligue Régionale contre le Cancer - Comité Gironde (n°267317 to S.M.). A.R.E. was recipient of a PhD fellowship from the Agence Nationale des Bourses du Gabon (ANBG).

## Supplementary data

**Figure S1.**
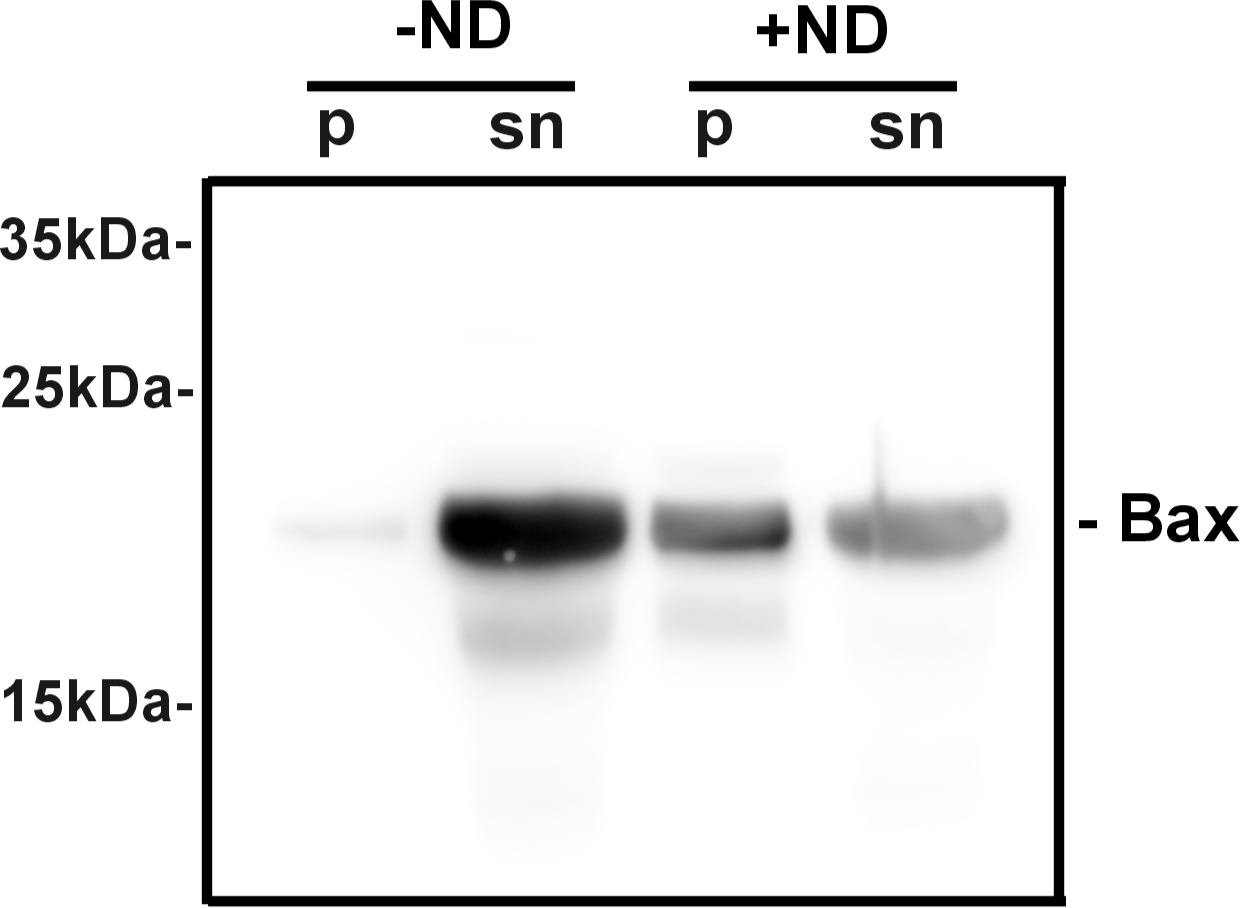
Precipitation of Bax induced by nanodiscs during cell-free synthesis. BaxWT was synthesized in the absence (-ND) or in the presence of 3.5µM nanodiscs (+ND). After the synthesis, the reaction mix was weighted and centrifuged at 20,000 x g for 15 minutes. The pellet was resuspended in the same volume of buffer B (25mM Hepes/Na pH 7.4, 200mM NaCl, 0.1mM EDTA). Aliquots from both the pellet (p) and supernatant (sn) were mixed with an equal volume of Laemmli buffer 2X, and analyzed by SDS-PAGE and western blot against Bax.

**Figure S2.**
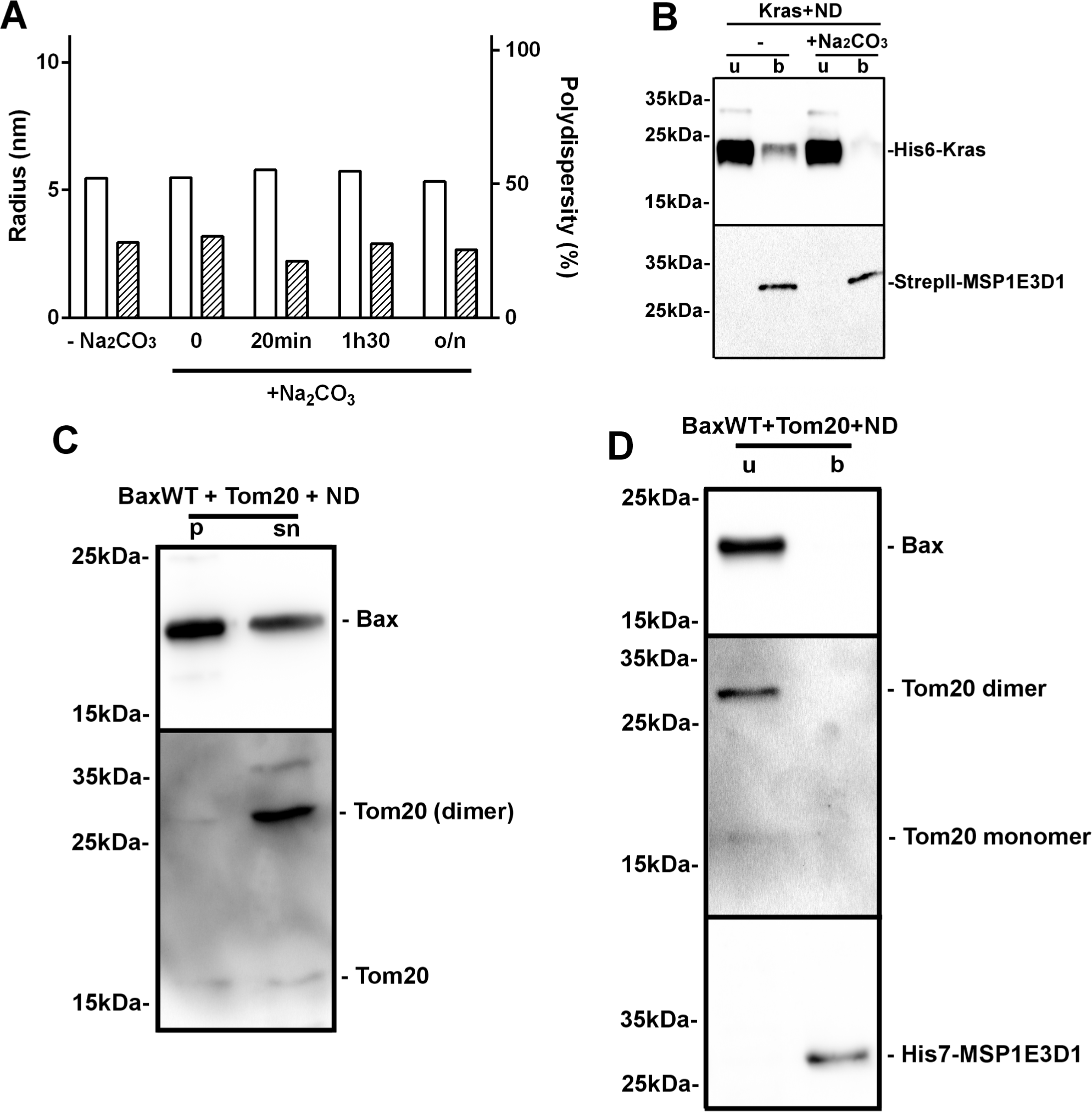
Controls of the effect of alkaline Sodium Carbonate treatment on nanodiscs and of the specificity of the effect of Tom22. **(A)** Nanodiscs were incubated for the indicated times in the presence of 0.1M Sodium Carbonate (pH 10.0). Radius and polydispersity were measured by DLS (3 measurements of 10 acquisitions each), showing that the integrity of nanodiscs was not affected by the treatment. **(B)** His6-tagged human Kras4b was synthesized in cell-free in the presence of StrepII-MSP1E3D1 nanodiscs. Nanodiscs were purified on StrepII-Tactin, then incubated for 15 minutes in the absence or in the presence of 0.1M Sodium Carbonate (pH 10.0). They were then re-purified on StrepII-Tactin, and the unbound and bound fractions were analyzed by western-blots. A minor fraction of Kras4b remained bound to nanodiscs, but was removed after alkaline treatment. **(C,D)** Bax was synthesized alone or co-synthesized with Tom20 and analyzed as in Fig.1A and 1C showing that Tom20 did not prevent Bax precipitation and did not promote its insertion.

**Fig.S3:**
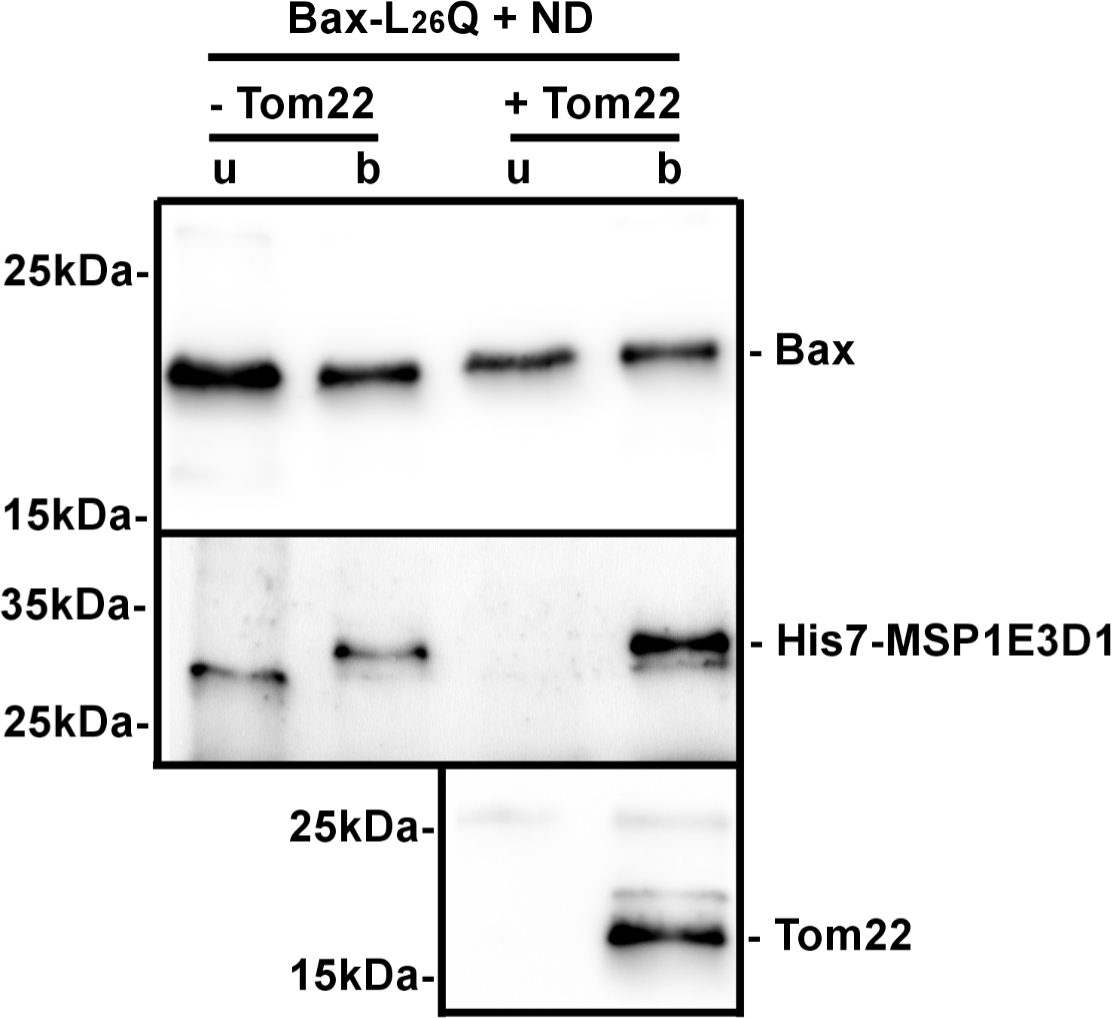
The mutation L^26^Q in the GALLL motif partially abolished Tom22-induced Bax insertion. Same experiment as in Fig.2H with BaxL^26^Q instead of BaxA^24^R.

**Figure S4.**
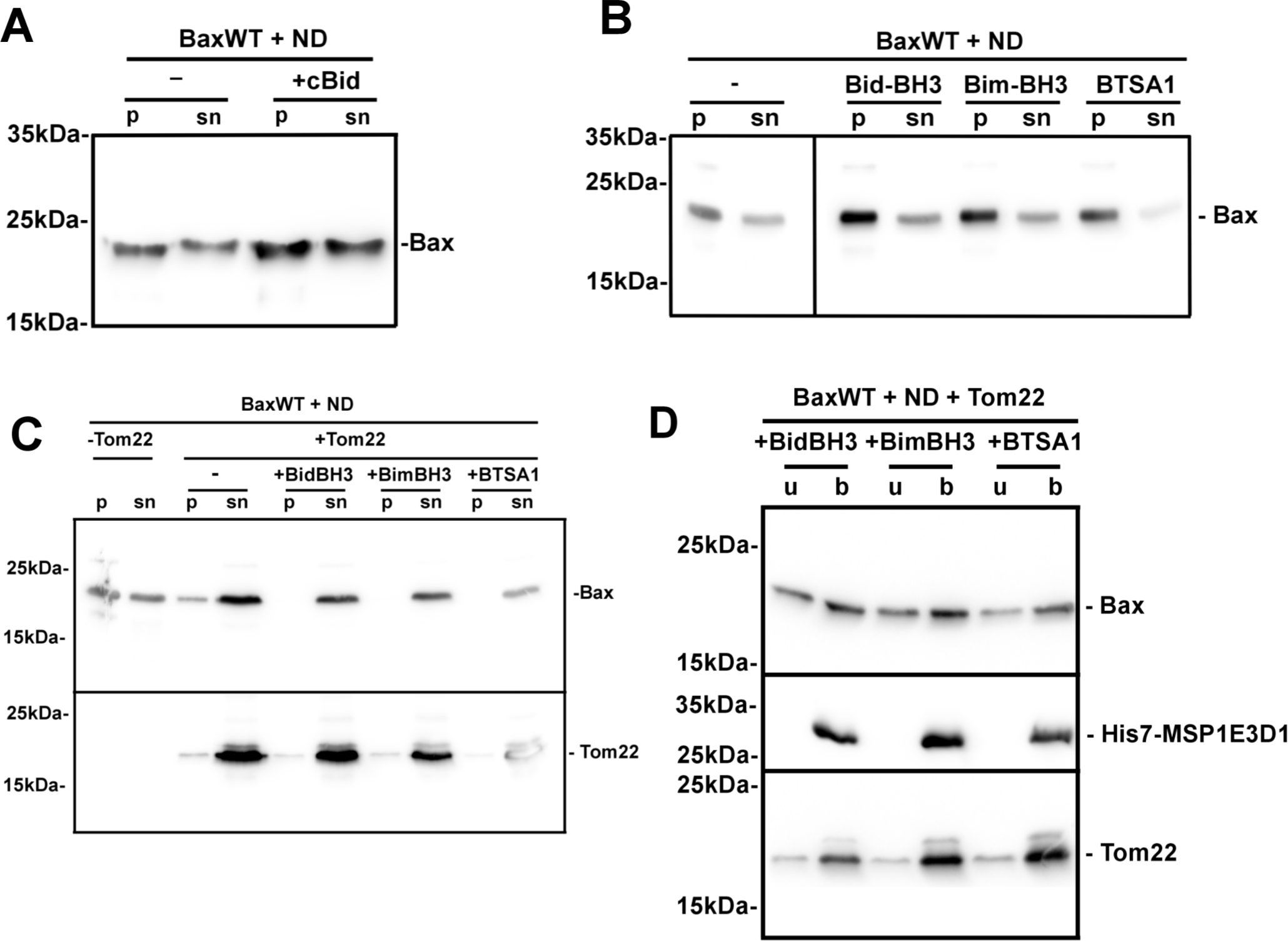
BH3-activators did not prevent Bax precipitation in the presence of nanodiscs. **(A)** Same experiments as in Fig.1A (without Tom22), except that 10µg caspase-8-cleaved Bid (cBid) was included in the reaction mix. We determined that the maximal concentration of Bax produced in the cell-free system was 0.5 mg/mL, *i.e.* 50µg of Bax in the 100µL-reaction mix. We therefore set up a cBid to Bax ratio of ∼1 to 5 (considering that the sizes of the two proteins are close to each other) (Shamas-Din et al., 2015). **(B)** Same experiment as in Fig.1A (without Tom22) in the presence of 1µM Bim-BH3 (PEIWIAQELRRIGDEFNAYYA), Bid-BH3 (ESQEDIIRNIARHLAQVGDSMDRSIPPG) (Genscript), or BTSA1 (Medchem). The concentration refers to the whole mix (reaction mix + feeding mix) because the sizes of the three molecules are below the cut-off of the dialysis membrane. **(C)** Same experiments as in **(B)** in the presence of Tom22. **(D)** Same experiments as in Fig.3B in the presence of Tom22.

**Figure S5.**
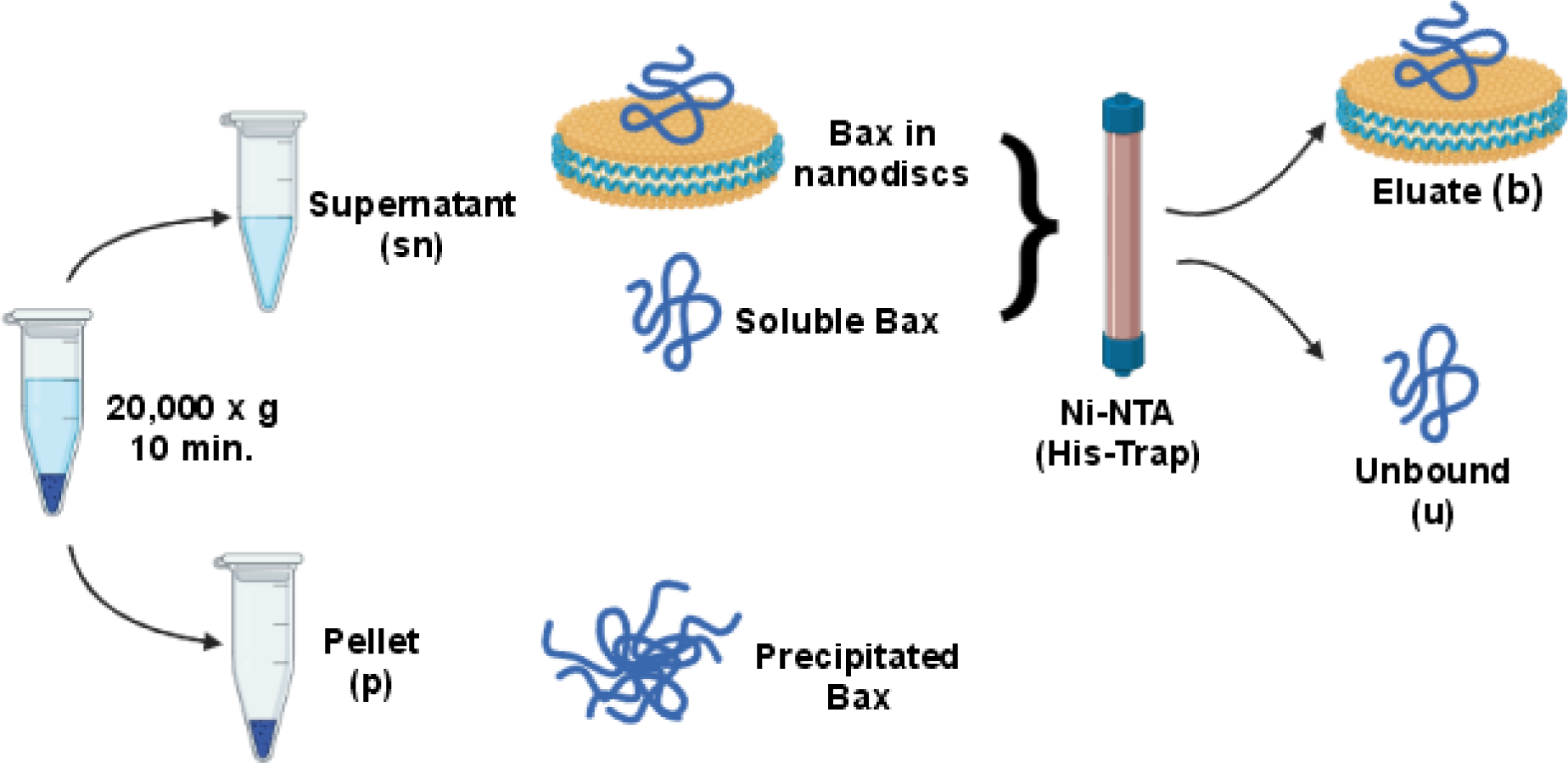
Schematic representation of the experimental flow. After cell-free synthesis, a 15-minutes, 20,000 x g centrifugation led to a pellet (p) of precipitated proteins and a supernatant (sn) containing soluble Bax and nanodiscs (empty or with inserted Bax). The supernatant was loaded on Ni-NTA (His-Trap). The flow-through contained soluble Bax unbound to nanodiscs (u). After an elution with 300mM imidazole, the eluate contained nanodiscs (empty or with inserted Bax) (b).

**Table S6:**
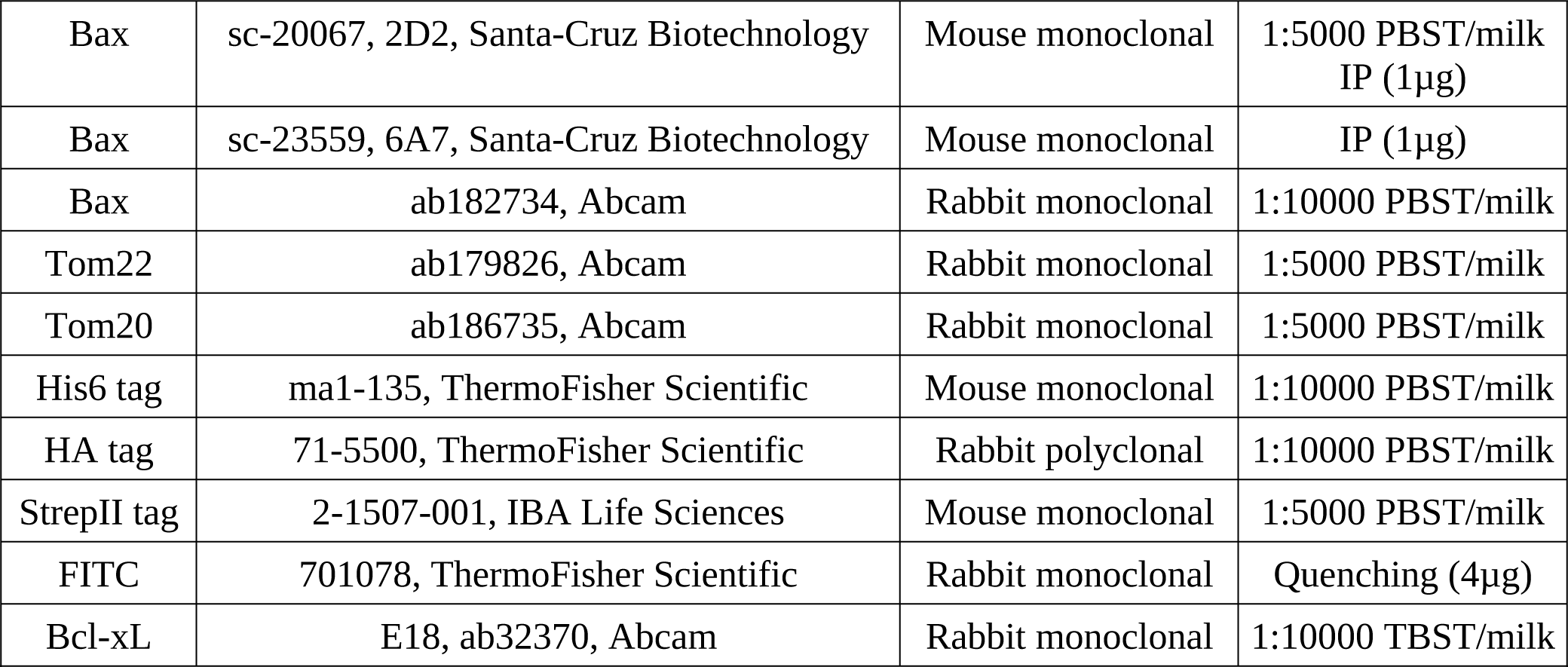
antibodies used in this study.

